# Midlife nonlinearities across scale in Biondi body amyloid formation in the human choroid plexus

**DOI:** 10.64898/2026.11.04.764372

**Authors:** Michael J. Neel, Natarsha Vukalovich, Parastou Porahang, Chenyu Zhao, William Huang, Sam Ghahremani, Andy Lin, Olivia Sandhu, Binh Nguyen, Kasra Pejhanfar, Erfan Zolfaghari, Eunice Shin, Carol A. Church, Joni L. Ricks-Oddie, John M. Ringman, Samuel Guzman, Peter D. Chang, Brett A. Johnson, Edwin S. Monuki

**Author notes:** Correspondence: Edwin S. Monuki.

## Abstract

Human choroid plexus epithelial cells (CPECs), which produce cerebrospinal fluid and form the blood-cerebrospinal fluid barrier, accumulate discrete intracellular amyloid inclusions called Biondi bodies (BBs). Using systematic manual and machine learning approaches on the choroid plexus glomus, we determined the percentages of CPECs containing thioflavin S-stained BBs as well as BB load and spatial distributions in whole slide images from 77 cases across the human lifespan. On average, BB prevalence was near zero until age 40, rose steeply in the 5th and 6th decades, and exceeded 50% after age 70. As prevalence plateaued, BB load continued to rise with age. Accelerated BB accumulation was seen in two cases of diabetic ketoacidosis, including a 16-year-old, while an 85-year-old with the relatively rare *APOE2/E2* genotype had fewer BBs than expected. BB prevalence in the choroid plexus glomus was not associated with Alzheimer’s disease, but was lower in females than males. Notably, spatial clustering of BBs was universal across cases while nearest-neighbor analyses suggested cell-to-cell, prion-like BB spread that can account for the nonlinearities seen at tissue and population scales. The abrupt, nonlinear BB accumulation during midlife coincides with or precedes other common aging-associated neuropathologies of the human brain, consistent with broad BB and ChP contributions to brain aging and disease.

## INTRODUCTION

The choroid plexus (ChP), a papillary secretory tissue within all four brain ventricles, produces and conditions the cerebrospinal fluid, which bathes the brain and spinal cord and contributes to central nervous system homeostasis [1–4]. ChP epithelial cells (CPECs) form the outer layer of the ChP and execute most of its functions, including forming the blood-cerebrospinal fluid barrier by way of their tight junctions [1–4]. With aging, the ChP undergoes numerous changes that likely threaten brain homeostasis [5–8]. Perhaps the best studied of these changes is the accumulation of intracellular amyloid deposits known as Biondi bodies (BBs) within the CPECs [8–20].

BBs were recognized in the 1930s as distinct features in silver stains of aged human ChP [9], and their amyloid nature was later made apparent through staining with Congo red [13] and thioflavin S [10–12], as well as through electron microscopy [14–17,20]. Along with the proteinaceous amyloid, most BBs contain an autofluorescent, lipofuscin-like component [15,20,21]. Consistent with this lipofuscin association, the lysosomal membrane protein TMEM106B recently has been identified as the amyloidogenic protein in BBs from numerous postmortem, neurodegenerative disease cases [20,22]. BBs are known both to increase in abundance during middle age and to become prevalent in most individuals by their 70s [10,13,16,19]. Increased prevalence in Alzheimer’s disease (AD) also has been reported [10,11], although this association has not been obvious in all studies [19]. In addition to AD, a case with multiple mitochondrial DNA deletions has been associated with accelerated BB formation [19].

Amyloid aggregates in general are known to exhibit cytotoxic effects by way of adsorbing other proteins and interfering with nuclear transport [23–26], and BBs in particular have been observed to distort or puncture CPEC membranes [11,17], suggesting that BBs may disrupt ChP function and thereby threaten brain homeostasis. Our laboratory has accumulated a large repository of ChP tissue collected at autopsy from standardized regions of human brain ventricles across the lifespan. As part of our systematic analyses of age-and disease-associated changes in the ChP, we stained sections of the ChP glomus (from the atria of the lateral ventricles) with thioflavin S to address BB abundance and spatial distributions using both manual and deep-learning approaches. Nearest-neighbor analysis revealed striking prion-like (cell-cell) spread of BBs, which plausibly underlies their spatial clustering within ChP and nonlinear midlife increase in prevalence within the population. In addition to age, we observed a sex difference and interesting outliers in BB prevalence, but found no association with sporadic AD, autosomal-dominant AD, or Down Syndrome in this study of the ChP glomus. Together with our accompanying paper, wherein we report that BBs exist in distinct, classifiable shapes that differ across individuals and in association with AD [21], these findings highlight the potential significance of BBs, CPECs, and ChP to human brain health and disease during aging.

## MATERIALS AND METHODS

### Human tissue

Human ChP autopsy samples were obtained from multiple sources including the UCI Medical Center (UCIMC) autopsy service, the UCI Alzheimer’s Disease Research Center (ADRC), the University of Southern California (USC) ADRC, and the Human Brain Collection Core (HBCC) at the National Institute of Mental Health (NIMH). From both UCI sources, ChP glomus was specifically collected (from the atria of the lateral ventricles where the frontal, temporal, and occipital horns meet) from brains fixed in either 10% formalin (UCIMC) or 4% paraformaldehyde (ADRC). From occasional UCI cases, ChP samples were also collected from the frontal and temporal horns. Postmortem intervals were typically lower for the ADRC compared to UCIMC cases (Table S1). ChP tissues from the HBCC were dissected from the lateral ventricle of frozen brain slices and shipped to UCI on dry ice, after which they were drop-fixed at room temperature in 10% formalin (Electron Microscopy Sciences, 15740-01) for 24 hours with gentle agitation. From the USC ADRC, one sample was provided as sections of formalin-fixed, paraffin-embedded (FFPE) tissue and the other was provided in PBS. All fixed, floating samples were processed by a UCI core facility (Experimental Tissue Resource) into FFPE blocks and sectioned into 5-µm thick sections.

### Thioflavin S staining and imaging

FFPE sections were deparaffinized using xylenes and rehydrated to 50% ethanol using a descending series of ethanol washes. Sections were then stained in 0.5% thioflavin S (ThS) (T1892-25G, Millipore-Sigma) for 10 minutes followed by two five-minute washes in 50% ethanol and three five-minute washes in phosphate buffered saline (PBS). After washing, sections were stained with 2 µg/ml Hoechst 33342 (#H1399, ThermoFisher) in PBS for 10 minutes and then washed two more times with PBS, five minutes each, followed by coverslipping with fluoromount G (#0100-01, SouthernBiotech).

ThS-stained FFPE sections were scanned into whole slide images at 20x (0.322 µm/pixel) using an Aperio Versa 2 scanner in the ADRC Neuropathology Core. Hoechst, thioflavin S, and autofluorescence were imaged using DAPI, FITC and TRITC/Texas Red filters, respectively. All imaging parameters including exposures were the same across samples.

### Manual Biondi body annotations

For manual analysis of BB prevalence, we employed a random image tile sampling methodology and stereology principles to estimate the percentage of ChP epithelial cells (CPECs) with BBs. First, ThS whole slide image files were converted from scn to svs format using Aperio ImageScope v12.4 software and then subdivided into 1024x1024 pixel (329.73 x 329.73 µm) image tiles. An example of an image tile is shown in Fig. S1a. Image tiles were presented in random order to annotators, who scored each CPEC containing a Hoechst-positive nucleus for presence or absence of any BB. After each image tile was annotated, cumulative BB prevalence was calculated. Starting with the tenth tile containing any CPECs, coefficients of error were calculated (standard deviations of prevalence across the last ten tiles containing CPECs expressed as a percentage of the current estimate). Once this coefficient of error remained below 5% across ten consecutive files, the case was judged as complete, and the data including prevalence values, CPEC annotations, and CPEC coordinates were exported. (By these criteria, the minimum number of tiles evaluated was 20.) If fewer than 2% of CPECs had BBs, annotation was halted at the completion of the last tile wherein 3500 CPECs or more were analyzed. Cell counts that were needed to meet these completion criteria ranged from 741 to 6358 with a mean of 2719 and standard deviation of 1133 based on 77 cases. Fig. S2 shows cumulative prevalence estimates for 3 different cases, indicating where the completion criteria were reached. Continued sampling beyond these criteria did not change the estimates appreciably.

Manual annotation was done in two phases. During the first phase (33 cases), ImageScope software was used for annotations, and progress towards criteria was calculated in spreadsheet software. During the latter phase, image tiles were generated as HDF5 files for use with a custom-designed software client to simplify blinded annotations, reduce sampling bias, and easily export data for DL model development. The client presented image tiles to annotators in two image windows representing different magnifications. The lower magnification window was used to locate CPECs and to use a mouse and keystrokes to annotate CPECs depending on whether they contained BBs. The client automatically signaled when completion criteria were met.

BBs were defined as ThS-positive structures within CPECs (Fig. S1b). Other green or ThS-positive structures often seen in tissue samples, such as fibrosis or elastin around large blood vessels (Fig. S1a), were ignored. Annotators included 9 undergrads and volunteers, as well as a senior project scientist (BAJ). Initial training on CPEC and BB classification was given by a board-certified neuropathologist with experience examining the ChP (ESM), followed by live demonstrations, video recordings, and regular feedback sessions with the project scientist. Annotations were conducted blind to all case and demographic information.

### Deep learning model development

Training, validation, and test datasets were developed using the annotations from the manual analysis of BBs, described above. For CPEC detection by the RetinaNet model, this included 8092, 1734, and 1734 images with annotated CPECs for training, validation, and test datasets, respectively. For BB classification, this included 24400, 6100, and 7625 CPEC images annotated for the presence of BBs for training, validation, and test datasets, respectively. For BB load, this included 5600, 1200, and 1200 images of individual, manually-segmented BBs for training, validation, test dataset, respectively.

To automate the quantification of BB prevalence, we developed a two-stage, deep learning (DL), convolutional neural network approach (CPEC detection followed by binary classification for BB presence). Two models were developed. First, a RetinaNet-based model [27] was trained to localize CPECs within an image, then a second ResNet-based model [28] was trained to classify CPECs with and without BBs. The first model adopted the RetinaNet architecture as described in Lin et al. [27] with modifications (Fig. S3). The model input image size was 256 x 256 pixels and the number of classes (K) and different anchor aspect ratios (A) were both reduced to 1. Because all images fed into the model had the same magnification and resolution, only the P3 feature map was passed to the regression and classification subnets. For the second model, a ResNet-based architecture [28] was used. This ResNet model had an input size of 64x64 and a fully-connected layer output of 2 (CPECs with and without BBs). To automate the quantification of BB load, we developed a U-Net like model [29] to segment regions of BBs. For BB load, a 4-layer U-Net like model was created with one output layer.

DL models were trained using Keras [30] and TensorFlow v2.5 [31] on a NVIDIA RTX 3090 graphics processing unit. For BB CPEC detection and classification, training of the two models was as follows: The RetinaNet model was trained using the Adam optimizer [32] with focal loss and Huber loss functions as provided in the jarvis-md python package (https://pypi.org/project/jarvis-md/). Before being passed into the model, images were first standardized to have a pixel mean of 0 and standard deviation of 1. The model was trained with mini-batches of 16 images with on-the-fly data augmentation including random contrast adjustment. During model inference, box predictions were pruned using a non-max suppression threshold of 0.3. The ResNet BB classifier model was trained using the Adam optimizer with a sparse categorical cross-entropy loss function. Similar to our RetinaNet training, images were standardized to have a pixel mean of 0 and standard deviation of 1 prior to being passed into the model. The model was trained with mini-batches of 16 images with on-the-fly data augmentation including random flipping, rotation, and contrast adjustment.

For BB segmentation, the model was trained using Adam optimizer with a sparse categorical cross-entropy loss function. Images were first standardized to have a pixel mean of 0 and standard deviation of 1 before being passed into the model. The models were trained with mini-batches of four 64x64 pixel (0.322 µm/pixel) images and incorporated on-the-fly image augmentation including random flipping, rotation, and contrast adjustment.

All relevant code regarding model architecture setup and training can be found in our GitHub repositories (https://github.com/mjneel/biondi & https://github.com/orgs/monukilab/repositories). Images and datasets are available upon request.

### Cluster analysis of Biondi bodies

Following annotations of every CPEC in a whole slide image, a local percentage of ChP cells with BBs was calculated based on cells within a 100-µm radius of each CPEC. For visualization, each CPEC was assigned a color based on the local percentage. Examples of raw and density-transformed data are shown in Figure S4. To test whether the spatial distribution of BBs differed from a random distribution, we generated Monte Carlo simulations using the same cell coordinates while randomly assigning BB presence based on the same overall prevalence value for a given section. We then used the Kolmogorov–Smirnov (KS) 2-sample test [33] to compare the distribution of measured local percentages and each of the 1000 Monte Carlo simulations to the combined distribution of all the simulations. A p-value was then generated by taking the number of simulations that had the same or higher d-value than the manual spatial distribution of BBs and dividing by the total number of simulations.

### Statistics

Statistical analysis was done under guidance of the UCI Center for Statistical Consulting. Spearman correlations as well as all spatial analyses were performed in python as described above. Analysis of covariance (ANCOVA) was conducted using SPSS version 28 software with robust error to account for heteroskedasticity in the data.

## RESULTS

### Development of a manual analysis pipeline for quantifying BBs

We first manually quantified BBs in human ChP autopsy samples. Due to the impracticality of quantifying entire tissue sections manually, we developed a sampling methodology and custom annotation client to estimate the percentage of CPECs with BBs (see Materials and Methods). This manual approach yielded average measurement standard deviations of 6.9% across different observers on independent sets of randomly sampled image tiles, and 7.3% when comparing multiple observers on the exact same images (Fig. 1). Thus, no additional variance was detected when tiles were sampled from different parts of the whole slide images. Despite the variance among annotators and sampled image tiles, the differences among the postmortem cases were statistically significant (ANOVAs: same image tiles, p < 0.001; different sampled tiles, p < 0.001).

**Figure 1:**
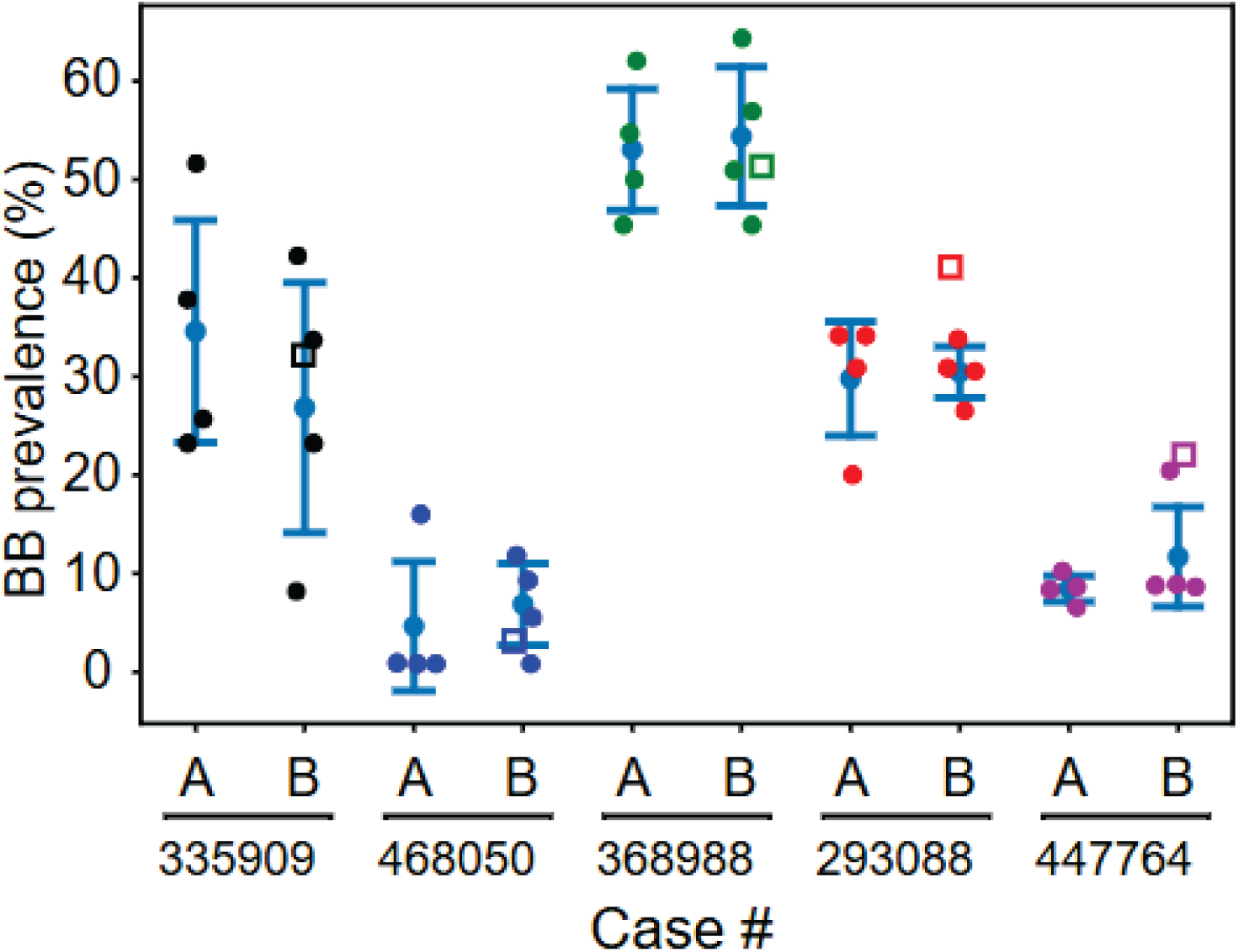
Interobserver differences are smaller than inter-sample differences. The point plot compares BB prevalence measurements across multiple different annotators and five different samples. Each color represents a different ChP sample. For each sample, BB prevalence measurements were made using either different random samples of image tiles for each annotator (**A; left**) or using the same set of image tiles across all observers (**B, right**). Circles represent data collected by annotators and hollow squares represent data collected using DL-models. Error bars denote standard deviations. Statistical tests confirmed that differences across cases were significant despite annotator variance for values in both **A** and **B**.

### Manual analysis of BB prevalence shows a nonlinear increase during midlife and a significant sex difference

BB prevalence was manually measured across 77 cases to explore factors that may impact BB abundance. Similar to previously published studies [10,19], the percentage of CPECs with BBs was near-zero below age 40, then increased dramatically in many, but not all individuals, during the 5th and 6th decades of life (Fig. 2). Over age 70, BB prevalence exceeded 50% of CPECs in most individuals, resulting in a highly sigmoidal (nonlinear) curve (Fig. 2), and this nonlinear increase was statistically significant (Table 1). To model the nonlinear increase, we fit the data to a Hill function (Fig. 2). The midlife period (operationally defined here as 50-65 years of age) displayed a high variability of BB prevalence across cases (1-70%).

**Figure 2:**
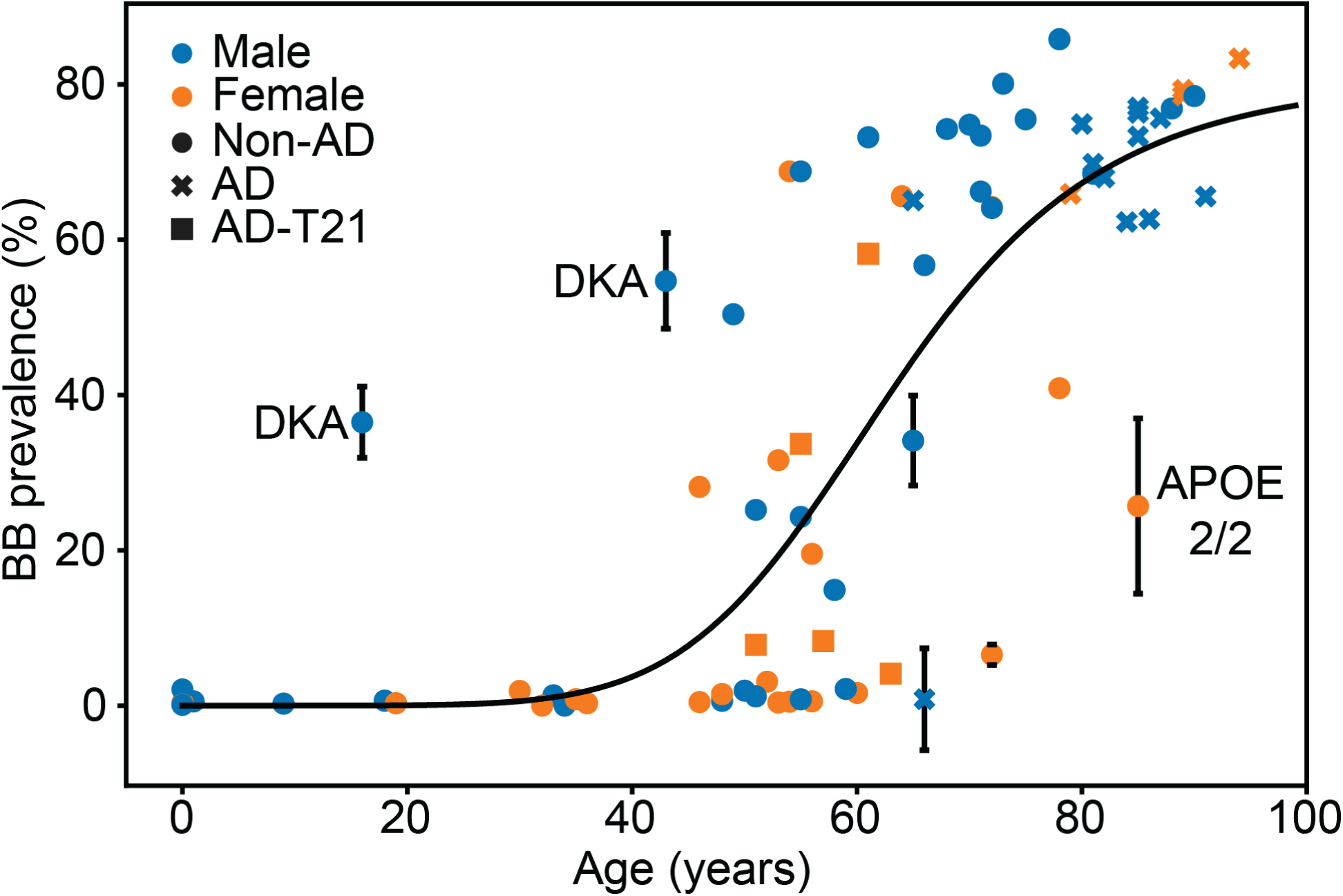
Manual analysis of 77 cases indicates that the percent of CPECs containing BBs increases nonlinearly with age, but not with AD. The curve is based on a Hill function fitted to the data (for constants, see Fig. 6). For cases used in the interobserver analysis (Fig. 1), the mean BB prevalence is reported along with error bars representing the standard deviation of independently sampled image tiles. “DKA” (diabetic ketoacidosis) and “APOE 2/2” (apolipoprotein E 2/2 genotype) label potential outlier cases.

**Table 1.** ANCOVA analyses of BB prevalence.

| Variable | p values (manual) |  | p values (DL model) |  |
|---|---|---|---|---|
|  | All cases | > 65 years old | All cases | > 65 years old |
| AD | N.A. | 0.136 | N.A. | 0.152 |
| Sex | 0.010* | 0.036* | 0.015* | 0.022* |
| Age | 0.606 | 0.361 | 0.525 | 0.341 |
| Age * Age | <0.001* | 0.431 | <0.001* | 0.401 |
| Age * AD | N.A. | 0.130 | N.A. | 0.136 |
| Tissue source | 0.930 | 0.290 | 0.504 | 0.120 |

Statistical analysis revealed a significant relationship with sex not reported in prior studies, with females having fewer BBs than males. This effect was still present when outlier cases were excluded from analysis (see below for outlier cases). No significant difference was seen with regard to the two main tissue cohorts (UCIMC autopsy service and UCI ADRC) used in the study.

### No systematic difference in BB prevalence across different ventricle locations

We analyzed twelve cases in our repository for which ChP had been collected from the frontal and/or temporal locations of the lateral ventricle in addition to the atrium location used for the bulk of our analyses. We detected no systematic difference in BB prevalence across the ventricle locations (two-tailed, paired t-tests: atrium versus frontal, p = 0.98; atrium versus temporal, p = 0.66) (Fig. S5, Table S1).

### BB prevalence was not affected by sporadic AD, Down syndrome, or PSEN1 mutation

Given previous reports of an association between BB percentages and AD [10,11], we tested for an AD relationship in our study. In contrast to prior reports, no relationship was found with AD status in individuals greater than 65 years of age (Table 1). Additionally, elevated levels of BBs were not detected in five cases with Down Syndrome (DS) and AD, which in DS is associated with overproduction of Aβ [33, 34], nor were elevated levels of BBs detected in either of two familial AD cases associated with PSEN1 mutations, aged 39 and 40 (unknown location in lateral ventricle and temporal horn location of ChP, respectively).

### Diabetic ketoacidosis and APOE2/2 cases displayed uncharacteristic BB accumulation

We identified a number of unique cases that did not seem to fall within the trend of BB accumulation. One particularly striking case was a 16-year-old male (case 363485) with 36.5% BB prevalence. To our knowledge, BBs have not been reported in individuals this young. This case had undiagnosed, uncontrolled type I diabetes and diabetic ketoacidosis (DKA) at death. Another case, an alcoholic 43 year old male (case 368988), also had diabetes and DKA at death as well as a previous episode of DKA. These cases suggest that uncontrolled blood sugar may promote BB formation.

Additional cases in our ChP repository were noted as having diabetes, including a 13-year old who died with DKA, but did not appear as outliers with respect to BB percentage (Table S1). To investigate further the possible association between BBs and DKA, we acquired 16 ChP autopsy samples (unknown lateral ventricle locations) from the Human Brain Collection Core (HBCC) at the National Institute of Mental Health (NIMH), half of which had DKA at time of death. These individuals were between 17 and 55 years old, and only the 55-year control case showed appreciable BB accumulation (Table S1). ANCOVA analysis of these cases failed to find a significant relationship between BB prevalence and DKA at time of death.

We also noted an 85 year old female (case 335909) with less than half the BB prevalence of other cases of similar age.This individual was the only case in our repository known to possess an apolipoprotein E (APOE) 2/2 genotype, which provides strong protection against AD [35].

### Manual analysis reveals spatial clustering of BBs

We noticed that the abundance of BBs seemed to vary across locations in some tissue sections. For three such cases, including the 16-year old diabetic and the 85-year old APOE 2/2 case, as well as a case that qualitatively showed extreme spatial heterogeneity (case 499605), we mapped CPECs with and without BBs across entire whole slide images, requiring manual annotation of ∼40,000-160,000 cells per whole slide image. We then plotted the positions of all annotated CPECs, and we color-coded points based on the local (100-μm radius) density of BBs. Initial qualitative observations revealed regional differences in BB density across all three cases (Fig. 3a,c,e). In case 499605 and the 16 year-old diabetic case (363485), clusters were relatively large with striking contrast between clusters of high versus low BB prevalence. In the APOE 2/2 case (335909), clusters appeared smaller with less contrast between high and low prevalence regions. Long stretches of clustered BBs were sometimes separated from nearby entirely devoid regions by small areas of stroma (Fig. S6).

**Figure 3:**
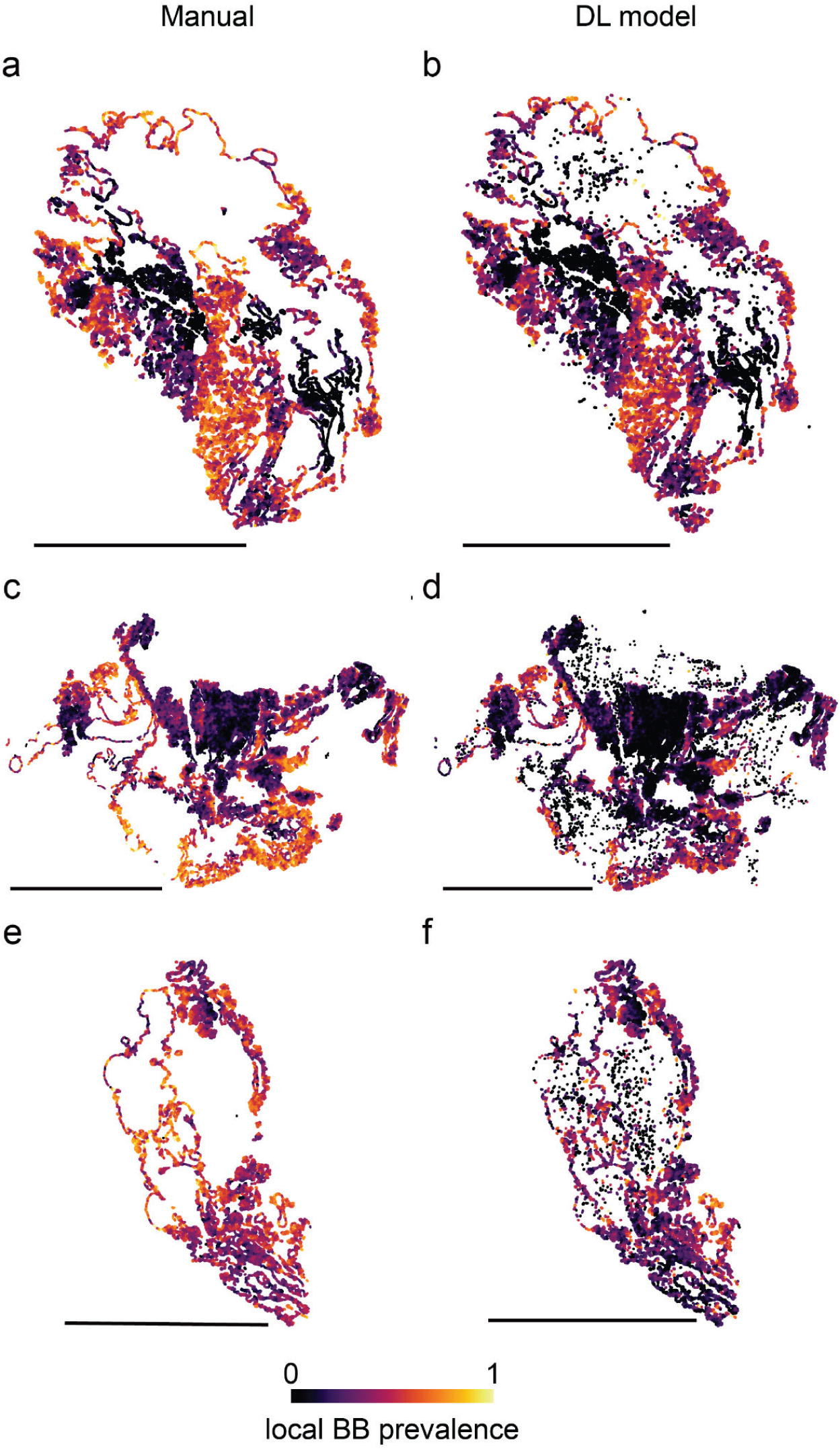
BBs are spatially clustered within the ChP. Manual and deep-learning model approaches were used to annotate CPECs across whole slide images of three cases for the presence or absence of BBs (**a,b:** case 363485; **c,d**: case 499605; **e,f**: case 335909). Each point represents a detected CPEC and is color coded to represent the density of CPECs with BBs within 100 μm of that point. Manual and DL approaches gave overall similar patterns of BB clustering, although DL models falsely reported several CPECs in large stromal compartments and failed to detect some CPECs that were annotated manually. Scale bars indicate 1 cm.

Histograms of local BB densities for the three cases (Fig. 4a,c,e, blue bars) revealed different types of distributions (unimodal for case 335909, skewed for case 499605, and bimodal for case 363485). To quantify the spatial clustering of BBs, we compared the observed distribution of local BB densities with 1000 Monte Carlo simulations of random distributions representing the same percentages of BBs (Fig. 4, average results shown as gray bars). In all three cases, the observed distributions were more different from the average random distribution than any of the 1000 simulations, indicating statistically significant spatial clustering of BBs, the magnitude of the difference being reflected in the Kolmogorov-Smirnov test d value (case 363485: d=0.24, p<0.001; case 499605: d=0.32, p<0.001, case 335909: d=0.21, p<0.001).

**Figure 4:**
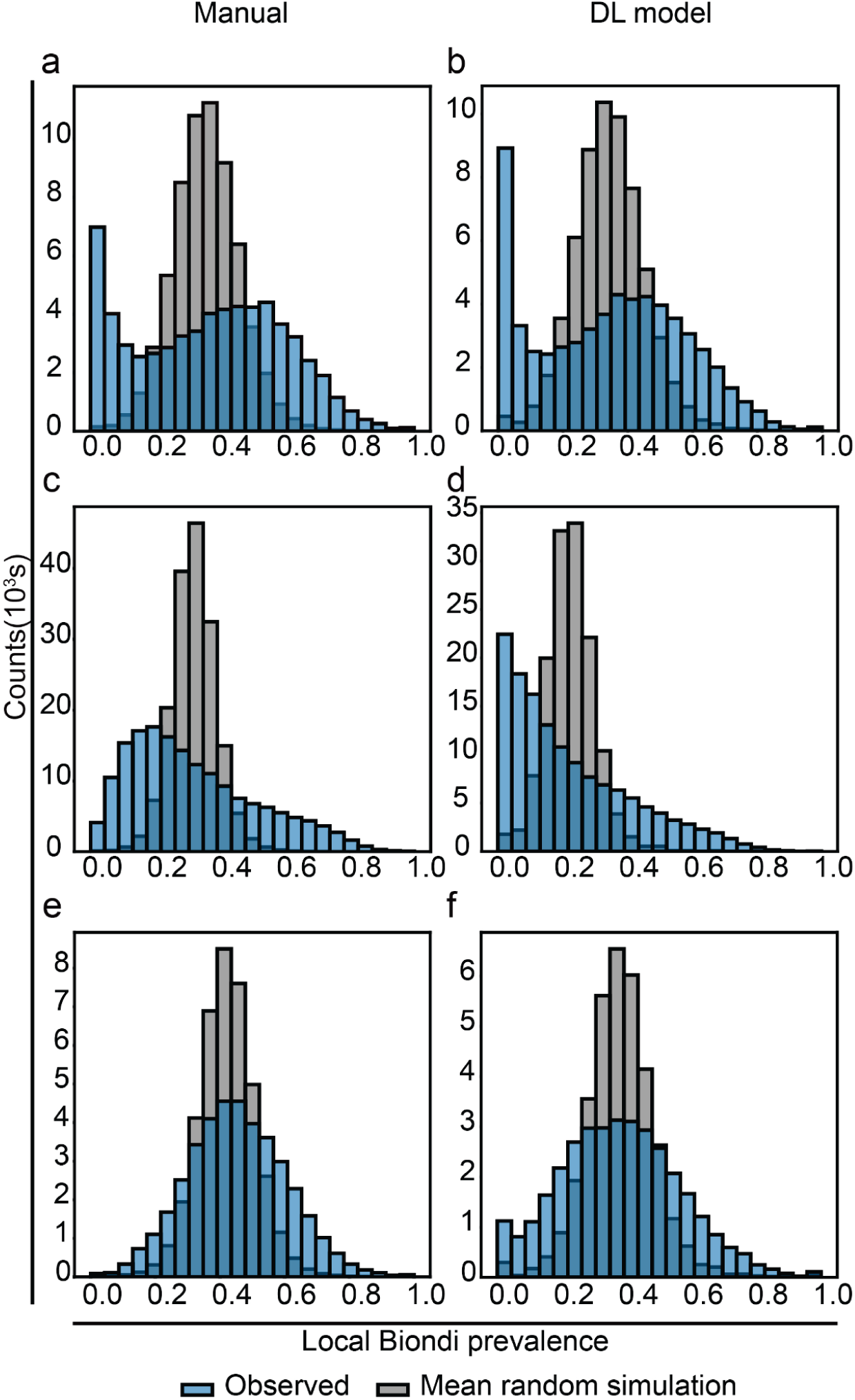
Histogram of local BB densities. Calculated for each cell shown in the scatterplots in Figure 3 (**blue**) compared to mean random distributions of BBs over 1000 Monte Carlo simulations (**gray**). Although the observed histograms varied among the cases (**a,b:** case 363485; **c,d**: case 499605; **e,f**: case 335909), they all differed significantly from random (p<0.001, Korsakov-Smirnov tests).

### Generation and use of deep-learning models to expand analysis of BBs

Given the striking spatial clustering of BBs seen in the three cases we analyzed manually, we sought to expand these analyses to other sections. Because the time and resources required to analyze a single whole slide image manually are high (∼40-80 person-hours of annotation per whole slide image), we chose to develop deep-learning (DL) models to automate the counting and categorization of CPECs and BBs. One of the biggest hurdles in the development of DL models is the generation of annotated image datasets for model training. We trained a RetinaNet-[27] and a ResNet-based [28] DL model to detect CPECs and classify CPECs based on BBs, respectively. In our case, the manual analysis of BBs generated ∼11500 images of ChP with annotated CPECs for ResNet training and ∼38000 images of individual CPECs labeled based on BB presence for RetinaNet training, which were adequate to train, validate, and test DL models. After training and hyperparameter optimizations the RetinaNet-based model achieved a median intersection over union (IOU) of 0.67 for boxes drawn around CPECs by the model compared to manual boxes, and the ResNet-based model classified CPECs with BBs with an accuracy, sensitivity, precision, and area under the receiver operating characteristic curve of 0.84, 0.70, 0.84, and 0.96. See Materials and Methods for additional details.

### DL models replicate results from manual analysis

To further validate our DL-approach, we compared our manual analysis of BB prevalence to our DL-model results. To quantify the correlation between prevalence measurements from manual and model data, we looked at model measurements based on the exact same image tiles used in the manual analysis (Spearman r = 0.92) as well as DL analyses of entire whole slide images (r = 0.95, Fig. 5a). Bland-Altman plot analysis [36] showed agreement between the two methods, with most measurements falling within the established limits of agreement (Fig. 5b). Open squares in Figure 1 show the results of the DL model annotations along with those of the human annotators from our interobserver analysis,

**Figure 5:**
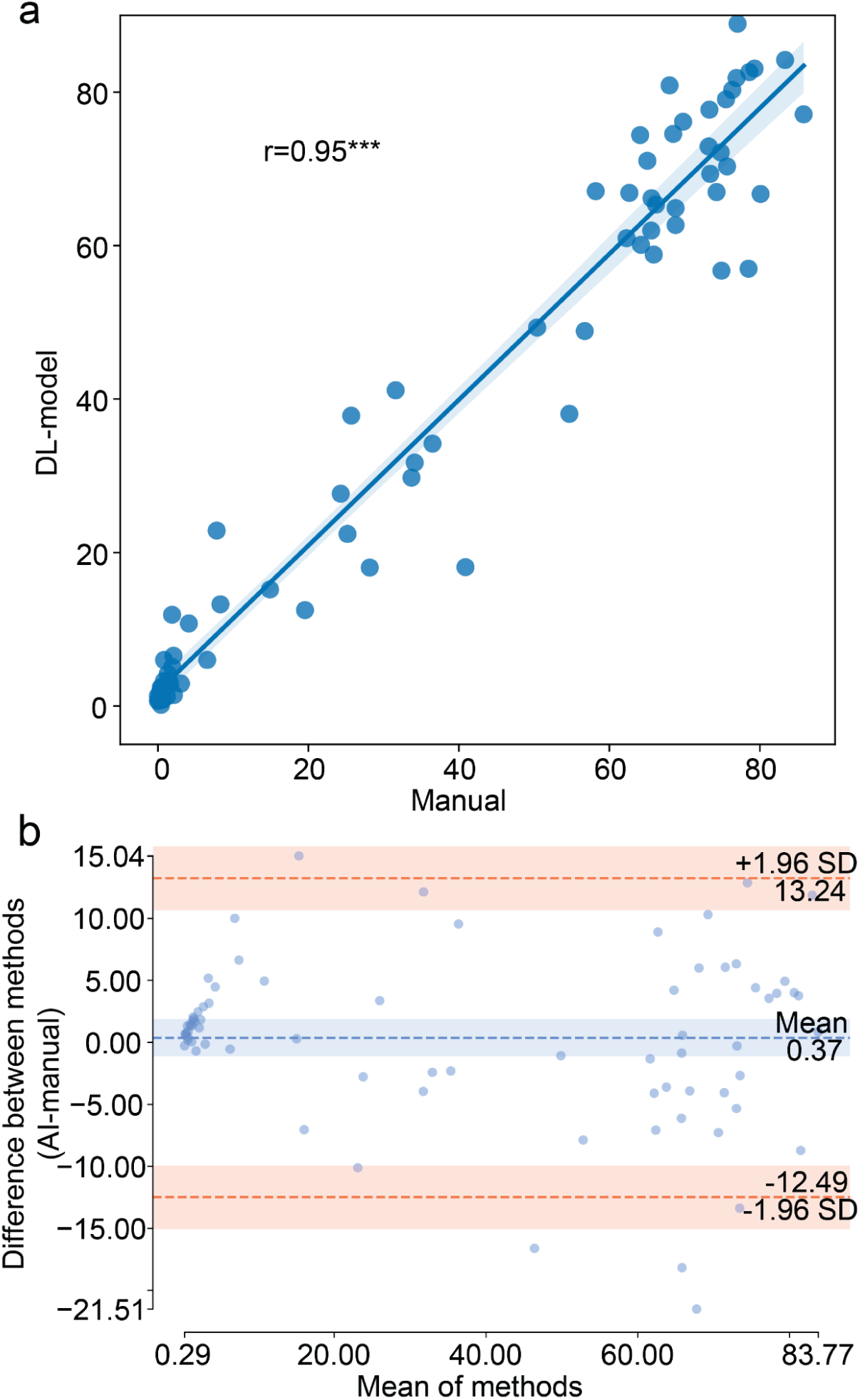
DL-models perform similarly to manual methods. **a**) Correlation analysis comparing manually acquired data to DL-model generated data. Linear regression is shown with 95% confidence intervals along with Spearman correlation coefficient (***, p < 0.001). **b**) Bland-Altman plot comparing manual and DL-based methods. Shaded regions represent the 95% confidence intervals of the mean and standard deviations of the differences between the methods (SD).

When analyzing the same 77 cases from the manual analysis, the model-generated data displayed a sigmoidal relationship between BB prevalence and age similar to our manual data (Fig. 6a). Indeed, the Hill functions fit to the two data sets were nearly superimposable (Fig. 6b), and the cases with uncharacteristic rates of BB accumulation in the manual data, including the pediatric DKA case and the APOE 2/2 case, also stood out in the model data. Statistical analysis (Table 1) had findings similar to the manual analysis, with significant associations between BB prevalence and both age and sex, and without significant association between BB prevalence and AD.

**Figure 6:**
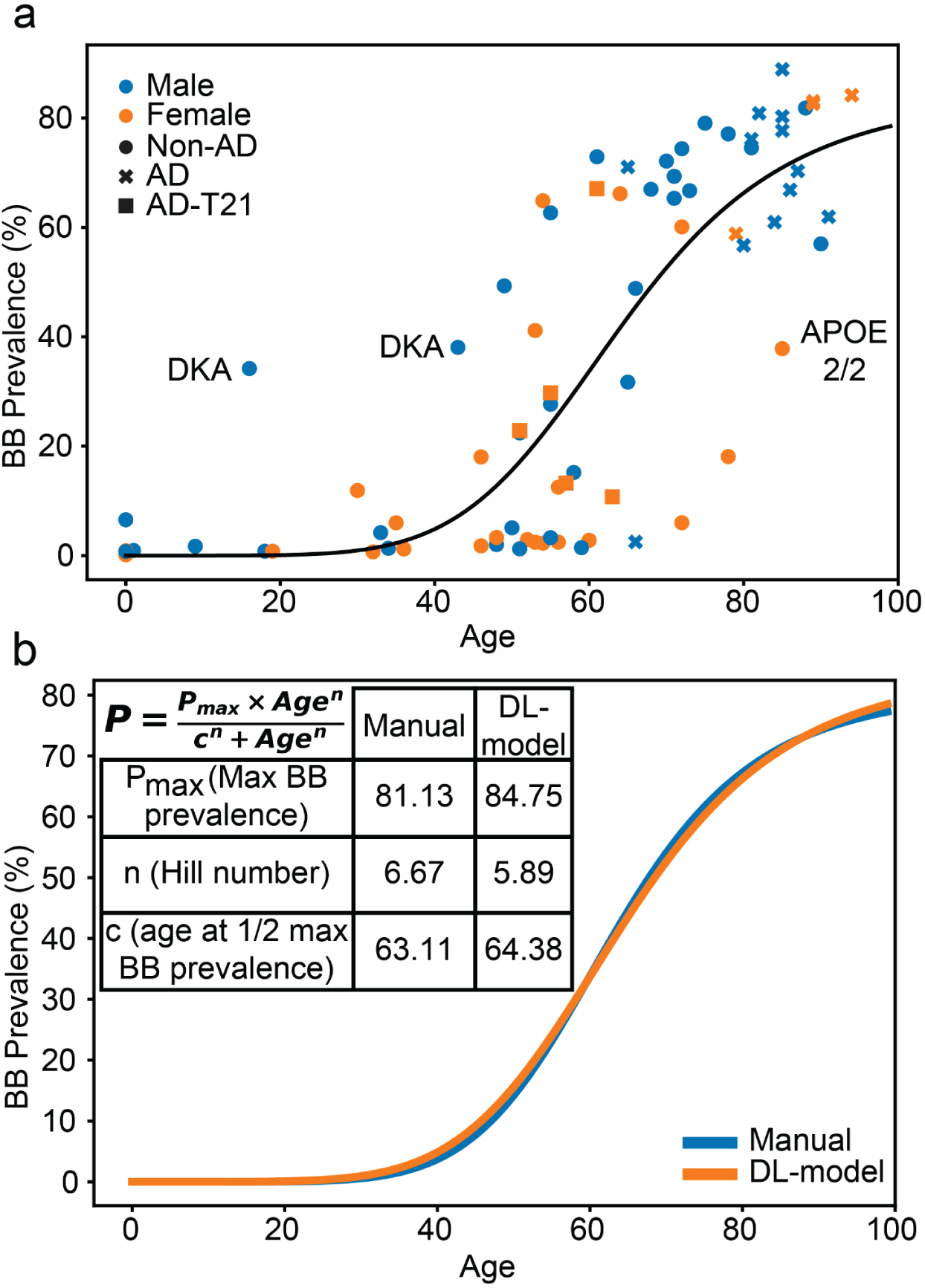
DL models recapitulate manually-acquired data. **a**) Scatterplot showing DL-generated BB prevalence data from the same 77 individuals in the manual analysis (Fig. 2). The curve is based on a Hill function fitted to the data. Note that BB prevalence increases nonlinearly with age starting around 50 years of age, similar to the manual analysis. The same cases noted as potential outliers in the manual analysis also fall outside the range of other cases of similar age in the DL analysis. **b**) Combined graph of Hill function curves from the manual-and model-generated BB prevalence data. Note the similarity between the two curves and Hill coefficients.

### DL models reveal spatial clustering of BBs in all cases

Having established that the DL models can replicate manual analyses of BB prevalence, we used them to analyze spatial clustering. The graphs of local density for the three cases annotated manually are shown alongside the DL model versions in Figure 3.The models show very similar clustering of BBs, although the DL model falsely annotated cells in the stroma as CPECs and did not detect as many CPECs as were annotated manually, which likely contributed to both a lower contrast in the plots of local BB density (Fig. 3) and a shift in local density histograms to lower values of prevalence (Fig. 4).

The DL models revealed significant clustering (p < 0.001) in all 77 cases analyzed when compared to Monte Carlo simulations of random distributions followed by Kolmogorov-Smirnov tests. The degree of clustering as measured by the Kolmogorov-Smirnov D value increased with age, but was highest at intermediate levels of BB prevalence (Fig. S7), which would be predicted given that extremes of high or low BB prevalence would necessarily be more homogeneous and have less opportunity to display clustering. However, clustering could clearly be seen even at these extremes. Samples with overall low levels of BBs had small regions that were highly affected, whereas samples highly impacted by BBs had small regions where CPECs were unaffected (Fig. 7).

**Figure 7:**
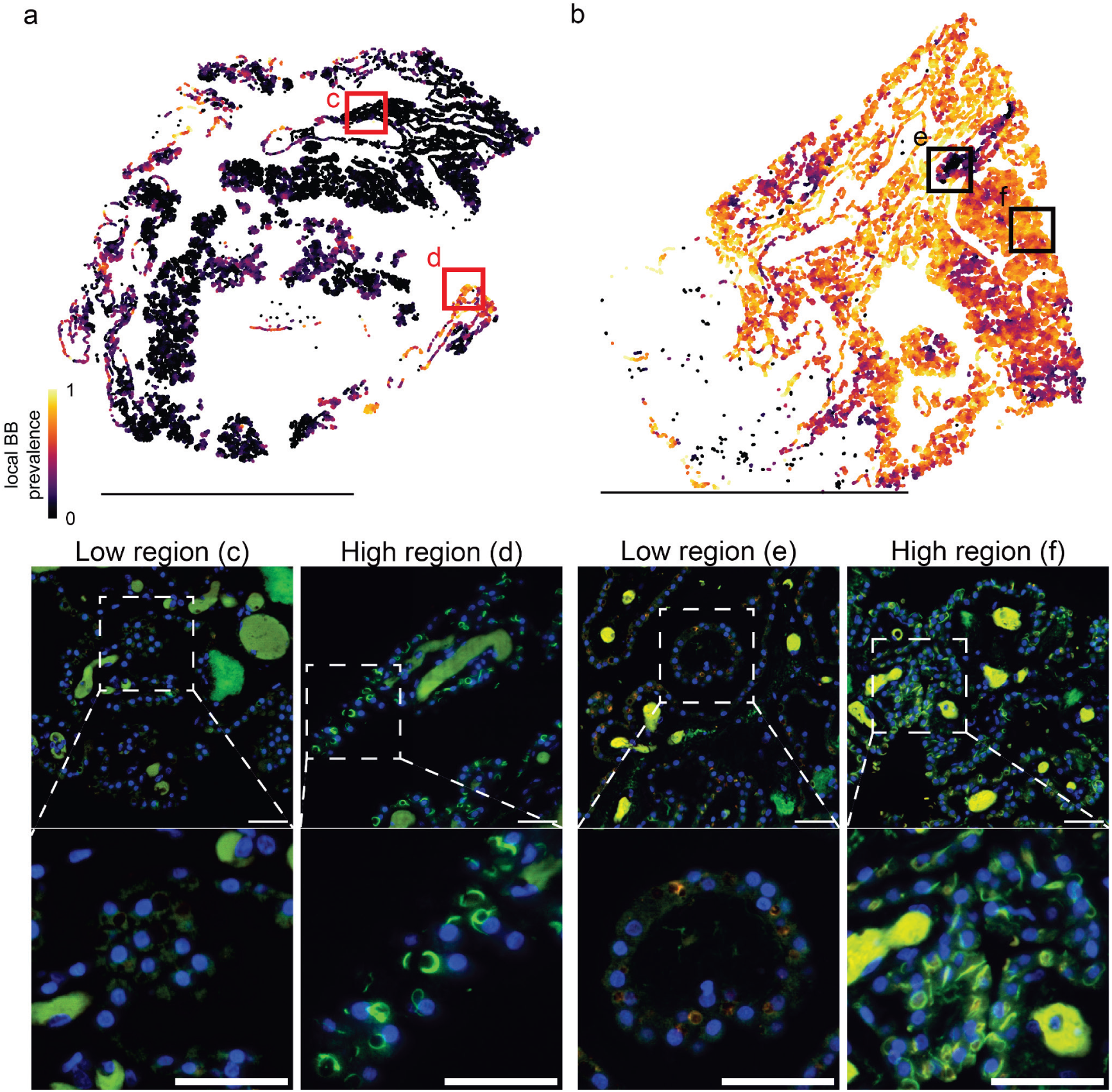
BBs cluster spatially regardless of prevalence. **a,b**) Scatter plots showing the location of annotated CPECs in whole slide images. Each point is color coded to represent the density of CPECs with BBs within a 100 μm radius. Scale bars = 1 cm. **a**) A case (#185324) with low BB prevalence (14.9%; manual) shows small clusters of high BB density. **b**) A case (#553262) with high prevalence (73.2%, manual) shows small clusters of low BB density. **c-f**) ThS images of boxed regions above (the red squares in **a** and **b** are centered on the regions depicted in c-f, but cover a larger area. Scale bars = 50 microns.

We also explored spatial clustering of BBs at a cell-by-cell level. Using our model-generated data, we calculated the probabilities of BB+ and BB-CPECs neighboring a BB+ CPEC in all 77 cases. We found that BB+ CPECs are significantly more likely than BB-CPECs within the same tissue to have adjacent BB+ CPECs (paired t-test; p=1.8 x 10^-33^) (Fig. 8, left). When looking at these probabilities in relation to BB prevalence measurements, the probability of BB-CPECs having a neighboring BB+ CPEC increases linearly with prevalence (Pearson r = 0.99, p<0.001), whereas the probability for BB+ CPECs exhibits a logarithmic-like increase with prevalence (Fig. 8, right). While BB+ CPECs were more likely to neighbor BB+ CPECs in every case quantified in this study, this effect appears strongest around intermediate prevalence levels (20-40%).

**Figure 8:**
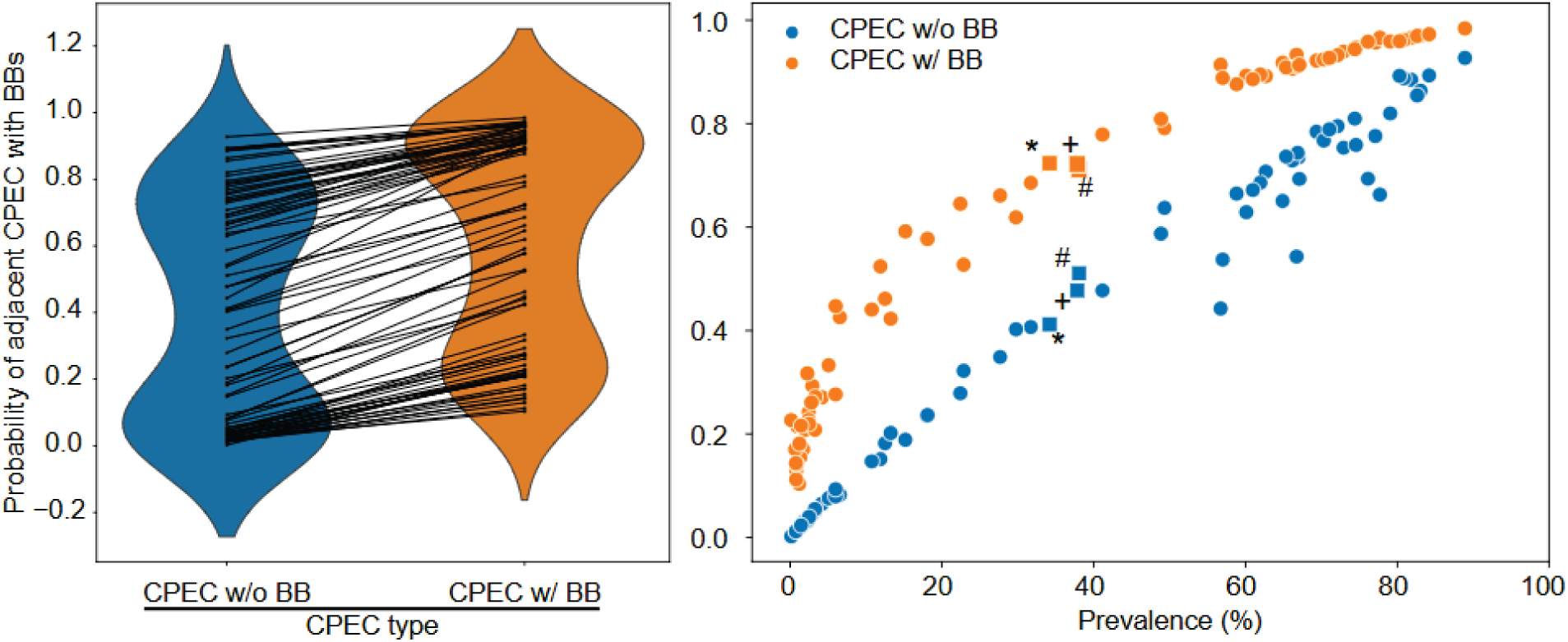
BB+ CPECs are more likely to neighbor other BB+ CPECs. (Left) Violin plots comparing the probability of a BB+ CPEC neighboring a CPEC with or without a BB across the different cases. Lines between the violin plots connect data from the same case. **(Right)** Scatterplot showing the probabilities previously mentioned as a function of BB prevalence.

We were concerned that BB classification of immediately adjacent CPECs could be affected by an overlap between DL model-generated bounding boxes (BBs in an affected CPEC might fall into the box for an adjacent, unaffected CPEC). To address this alternative explanation, we also tested the probabilities of BB+ and BB-CPECs one further CPEC removed from a BB+ CPEC in the same 77 cases. This analysis yielded the same conclusion that the probability of finding a BB+ CPEC near another BB+ CPEC was higher than finding a BB-CPEC (paired t-test, p= 2.0 x 10^-23^) (Fig. S8).

For each of the 77 cases, we correlated the local density of BBs at each position with the local density of CPECs. A one-sample t-test of the Spearman correlation coefficients revealed that the correlations were significantly different from zero (t = 5.92, p = 9 x 10^-8^, mean r = 0.086), reflecting higher densities of BBs where there were higher densities of CPECs.

### DL model load analysis

Because BB prevalence on its own does not fully reflect the abundance of BB amyloid material in the ChP (BBs could continue to increase in size or number in each CPEC), we used DL models to quantify BB load. To generate training, validation, and test datasets, we manually segmented BBs in 8000 CPEC images generated from our BB prevalence DL-model. These datasets were then used to train and validate a UNet-like DL-model [29] to segment BBs and generate a BB load measurement. On our test set, the resulting BB load model achieved a dice score of 0.57 (2 times the area of overlap between manual and DL model annotations/sum of the areas of the two annotations). We then used this model to quantify load across the 77 cases used in this study. BB load increased significantly with age (Table 2), and in an exponential fashion (Fig. 9A), without the plateau apparent in the prevalence curves.

**Figure 9:**
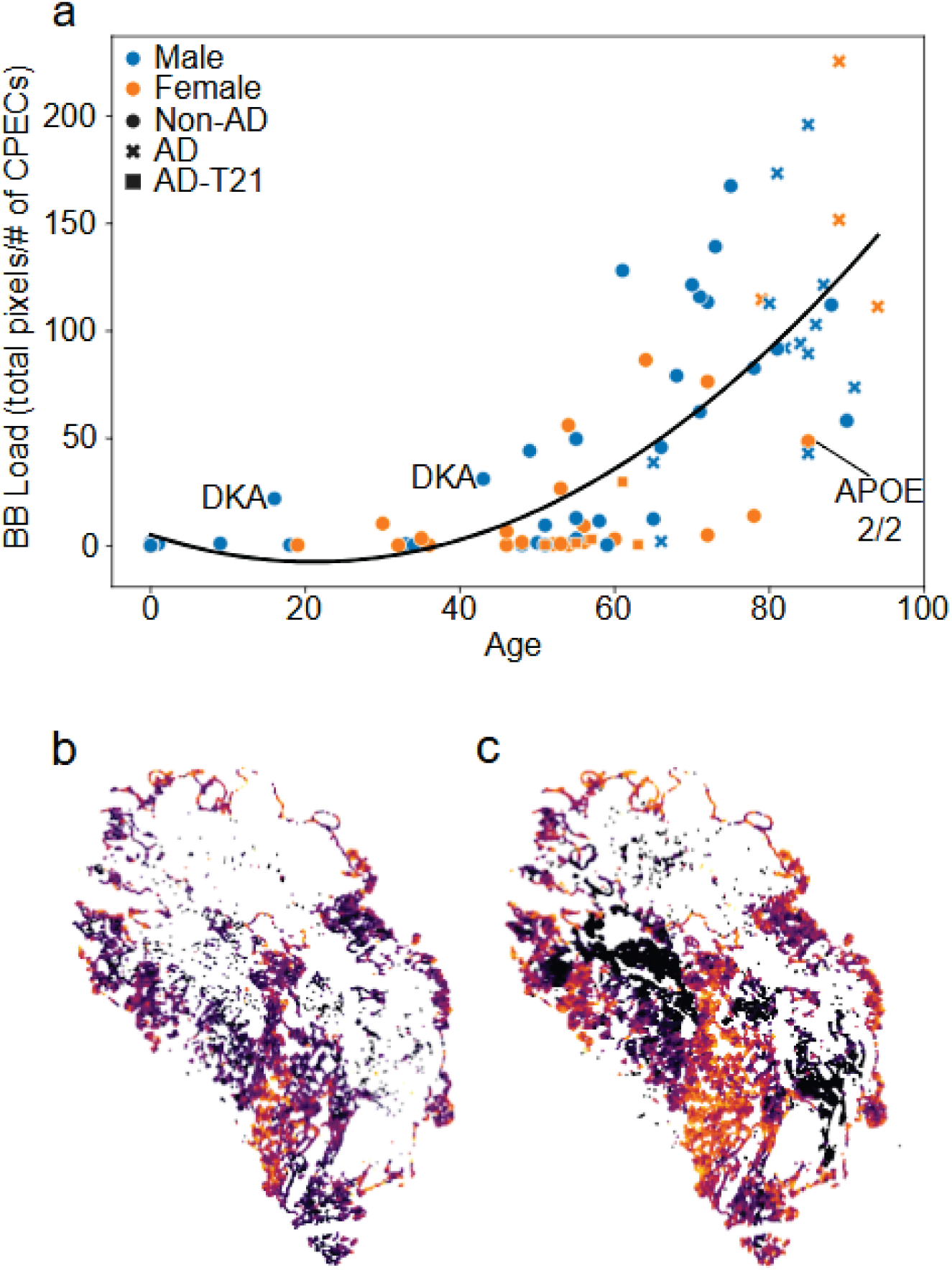
BB load increases with age and is higher in the center of a cluster. **a**) Scatterplot showing DL-generated BB load measurements as a function of age. The trendline is a 2nd order polynomial regression, showing that BBs continue to expand with increasing age. **b**) Heatmap showing the spatial distribution of local BB load density for case 363485. **c**) Heatmap showing the DL-generated local BB prevalence density (the same as in.Fig.3b). Note how BB load (**b**) is higher towards the center of the prevalence cluster (**c**).

**Table 2.** ANCOVA analyses of BB load.

| Variable | p values (DL model) |
|---|---|
|  | All cases |
| Sex | 0.060 |
| Age | 0.014* |
| Age * Age | <0.001* |
| Tissue source | 0.236 |

Lastly, we addressed whether BB load might vary across a cluster of BBs. For this analysis, we chose the 16-year old diabetic case possessing a large, distinct BB cluster (case 363485), and we generated a heatmap based on the local load of BB+ CPECs (Fig. 9B). BB load appeared higher toward the center of the BB-prevalence based cluster (Fig. 9C).

## DISCUSSION

In this study, we used manual and deep learning AI approaches to characterize quantitatively an abrupt midlife increase in BB accumulation. Although we did not find an association between AD and the percentage of CPECs with BBs, we did find that females had a significantly slower accumulation of BBs. We identified two cases involving uncontrolled blood sugar that displayed a higher prevalence of BBs than expected for their young age, as well as an 85-year old case with the AD-protective APOE 2/2 genotype that showed a lower prevalence of BBs than expected. BBs were found to be spatially clustered in every ChP studied, suggesting the possibility of a prion-like spread.

### Comparison to prior studies

A late onset of BB formation was appreciated in early semi-quantitative studies of postmortem ChP [8,16,18], and our quantitative data showing an age-dependent, nonlinear, midlife increase in the percentage of CPECs containing BBs largely agree with prior quantitative studies, which are plotted along with our manually acquired results in Figure 10A [10,19]. All three studies showed a great deal of variability across individuals in the percent of CPECs affected over the middle-age range of about 50-65 years. It is unclear how much of this variability is biological and how much is technical. All of the data comes from analyses of single histological sections through tissue that we now know to contain a spatially clustered distribution of BBs. Therefore, sampling error due to clustering in the z-plane would contribute to the variability.

**Figure 10:**
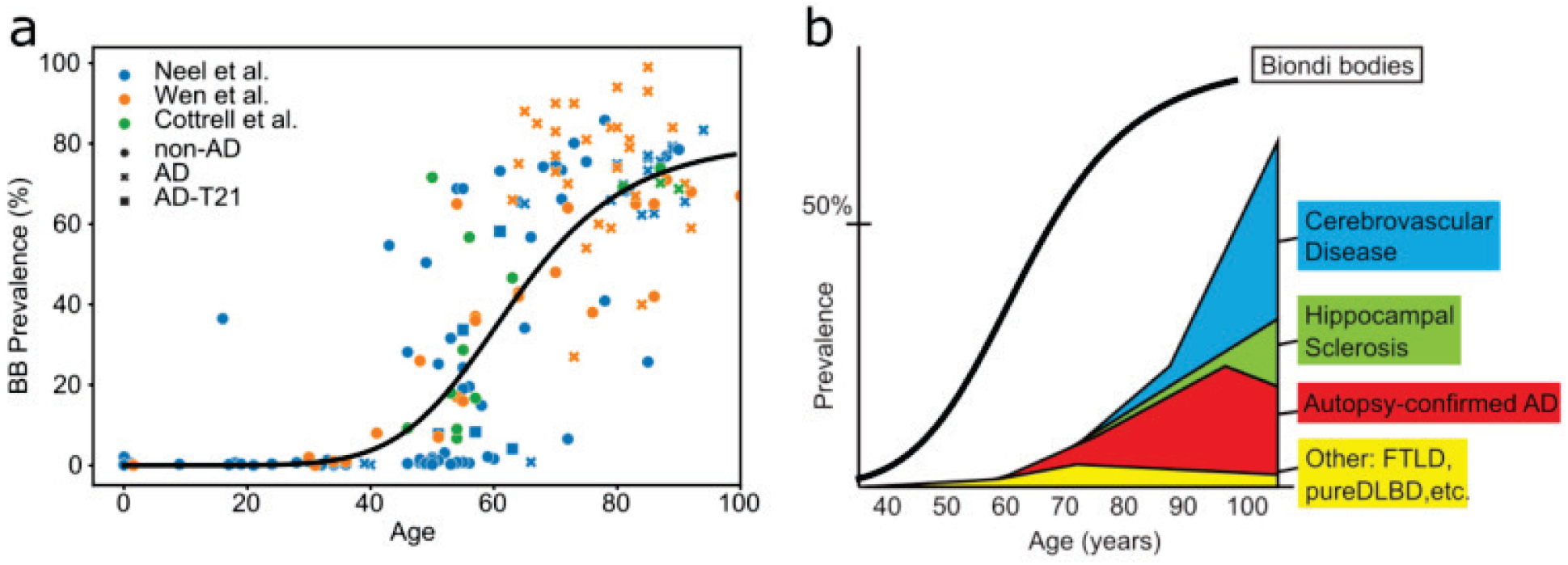
Comparisons of our results to prior studies. **a**) Manually acquired data from this study are plotted together with values from two other studies that quantified the percent of CPECs containing ThS-positive BBs [10,19] (Table S1). While in general agreement concerning the rapid rise of BBs around middle age, the Wen et al. [10], non-AD values fall below the range of values from the other two studies. The multiple mitochondrial DNA deletion case from the Cottrell et al. study [19] continues to show a greater accumulation of BBs than other, similarly aged cases when plotted together with our larger dataset. **b**) The Hill function fit to the DL model-acquired data from this study is plotted along with a diagram modified from Nelson et al. [65] illustrating the age-dependent accumulation of various neuropathological changes. BB accumulation precedes these signs of neurodegenerative disease. Note that “prevalence” of BBs indicates percentages of affected CPECs whereas prevalence of other pathologies indicates percentages of people afflicted.

In the Wen et al. study that reported an association of BBs with AD [10], values from older, non-AD individuals tended to fall below both our values and values from the Cottrell et al. study [19] within this age range. Statistical analysis of our data did not reveal an association with AD, in contrast to both Wen et al. [10] and a prior semi-quantitative study by Miklossy et al. [11]. We also did not detect increased BB accumulation in ChP from either Down Syndrome patients with AD or familial AD patients with presenilin 1 mutations, showing that fully developed AD neuropathology can be present with little to no accumulation of BBs.

The Miklossy et al. study finding increased BB in AD combined results from ependymal cells with results from CPECs [11]. Although these cell types frequently have been conflated and although the amyloids in the two cell types include TMEM106B [20,22], the two cell types differ greatly in developmental origins (CPECs arise early in brain development and directly from neuroepithelial stem cells with no glial intermediate [37–39] while ependymal cells arise later in development from radial glial cells [40]), their mRNA and protein expression are very different (scRNAseq results in clearly distinct clusters [41,42]), ependymal cells respond in an astrocyte-like manner to inflammation while CPECs do not [42], and ependymal cells are directly adjacent to brain parenchyma, while CPECs are separated from brain parenchyma by CSF. Given the marked differences between the cell types, it is possible that AD is more associated with changes in the ependymal cells than in the CPECs.

Another potential difference in the studies is that the ChP specimens used in our statistical analysis were obtained specifically from the atrium of the lateral ventricle, whereas the prior studies did not provide details regarding the region sampled. Because the ChP in ventricle locations other than the atrium can be closer to brain regions that are highly impacted in AD (such as the temporal horn of the lateral ventricle adjacent to the hippocampus), it is possible that our different results reflect a difference in the location of the tissue extracted from the brain. This may be especially important if BBs can be seeded by other amyloid proteins present in AD (see below). We conducted a limited study comparing different lateral ventricle locations from the same cases and did not see a systematic difference (Fig. S5), but this comparison lacked the power to address an interaction with AD.

Finally, there may have been differences in the criteria for defining BBs in the different studies. In the present study, we considered any CPEC containing a high-contrast, ThS-stained object to be BB-positive, independently of BB size. In our study of BB morphology, AD was associated with an increase in complex BBs and a decrease in clumps of multiple, small BBs [21]. These smaller BBs may have been overlooked in the prior studies in favor of the larger BBs. It should be noted, however, that we also did not detect any association between BB load and AD, an analysis that might have been sensitive to AD-associated increases in BB size..

While not replicating an AD difference, our study did find that female individuals have lower BB prevalence compared to males, which contrasts with a higher beta-amyloid and tau pathology burden in females [43–46]. Genes associated with various ChP functions are known to be modulated by sex hormones in animal models [47–49], which may be relevant to the observed sex difference in BB accumulation.

Our study included two diabetic cases with a BB abundance greater than would be expected given their age, and both of these individuals had DKA at the time of death. One individual, a 16-year-old who had undiagnosed (and therefore untreated) type 1 diabetes, is by far the youngest BB case so far reported. The other case was a 43-year old alcoholic with a DKA hospitalization one year earlier, suggesting a history of uncontrolled blood sugar. ChP specimens from other individuals who died with DKA did not provide further evidence for a relationship between BBs and DKA, but the duration of uncontrolled blood sugar in these other cases could not be determined. The rate of amyloid formation of some proteins is accelerated with the glycation that accompanies high blood sugar [50–55], and glycation can neutralize lysine side chains and possibly decrease repulsion between protein chains, thus promoting their aggregation [55]. Given the striking outlier status of the 16-year-old case, further attention should be paid to a possible relationship between high blood sugar and BB accumulation. However, this case also suffered comorbidities (asthma, hepatomegaly, and cardiomegaly with accelerated atherosclerosis) that may have contributed to accelerated BB accumulation.

When plotted alongside our data, the 50-year-old multiple mitochondrial DNA deletion case reported as a positive outlier by Cottrell et al. [19] continued to fall outside the range of other cases at that age (Fig. 10A), consistent with an acceleration of BB accumulation accompanying other metabolic disturbances.

An 85-year-old individual with the very rare APOE 2/2 genotype displayed fewer BBs than other, similarly aged cases. APOE gene polymorphisms are the strongest genetic association with sporadic, late-onset AD, with the APOE4 isoform greatly increasing AD risk and the APOE2 isoform greatly reducing risk compared to the more common APOE3 isoform, while both isoforms display both protective and risk associations with other disorders [56,57]. Because the APOE genotype has not been determined for the majority of cases in our dataset, further study will be needed to assess more thoroughly any general relationship between APOE genotype and BB accumulation.

### Biondi body load

In addition to expressing the accumulation of BBs in terms of the percentage of CPECs containing any evidence of a BB, we also developed a DL model-based method of assessing the overall BB load. This assessment is sensitive not only to the number of cells with BBs, but also to the number and size of BBs within each CPEC, which would likely be more predictive of changed CPEC function. When expressed as load, BBs were seen to increase exponentially starting around middle age rather than reaching the artificial plateau characteristic of the percentage plots. The percentage plots are necessarily asymptotic at 100%, and our additional criterion requiring the presence of a visible nucleus for annotation would lead to undercounting of the affected CPECs.

### Spatial clustering and potential prion-like spread of BBs

Rather than being scattered randomly across the ChP epithelium, CPECs containing BBs were found in spatial clusters in all 77 cases investigated, and the clustering was determined to be statistically significant by two different methods. Even the most heavily affected cases had islands of unaffected cells. One potential explanation for this clustering is prion-like spread, in which misfolded proteins in one cell cause normally folded proteins in other nearby cells to misfold. Many amyloids are known to exhibit prion-like spread and seeding of new amyloid aggregates, including beta amyloid, tau, and alpha-synuclein [58,59]. We found that BB load was higher in the center of a large cluster compared to the periphery, which is consistent with a spreading model in which originally affected cells would have additional and larger BBs, while the CPECs to which the BB amyloid has more recently spread would have lower loads. Our finding that BB density is correlated with CPEC density may also support a mechanism of prion-like spread.

We observed clusters involving affected CPECs near one another along the monolayer as well as separated by narrow channels of CSF space. On the other hand, we observed clusters of affected CPECs on one side of a thin strip of stroma while CPECs on the other side were unaffected. This type of distribution favors potential mechanisms of spread involving gap junctions between CPECs [60] and/or apocrine secretion into the CSF [61] rather than diffusion through stroma. We are particularly intrigued by the possibility that BBs could cross-seed or be cross-seeded by other types of amyloid species in the brain. Heterotypical amyloid cross-seeding has been observed for other amyloid proteins involved in neurodegenerative diseases [62,63].

The phenomenon of BB spread suggests two potentially distinct factors influencing the percent of CPECs bearing Biondi bodies, namely the rate of amyloid initiation and the rate of amyloid spread. The large clusters of BBs detected in the 16-year old, untreated type 1 diabetes case, who had abundant BBs despite his youth, together with the relatively simple morphologies of the BBs in this case [21], may be best explained by acceleration of prion-like spread.

### nonlinear, age-related accumulation of Biondi body amyloid

Many age-related transcriptomic changes occur in the ChP during middle-age, prompting one group to describe the 50’s as “a transition decade, in choroid plexus aging” [7]. The rapid accumulation of BBs between 50 and 65 years of age is consistent with this conception, and it also occurs during the second crest of nonlinear, age-related changes found in a recent multi-omic study [64]. The wave of changes in blood transcriptome and proteome occurring at around age 60 includes, but is not limited to, changes in carbohydrate metabolism, cardiovascular disease, immune function, and extracellular matrix [64].

As illustrated in Fig. 10B, BB accumulation precedes age-related neuropathological changes, such as cerebrovascular disease, hippocampal sclerosis, and various dementias [65]. Given that BBs likely impair CPEC function and may cross-seed brain amyloids, it seems possible that BB accumulation puts the central nervous system at risk for neurodegenerative disease (Fig. 10B).

## ACKNOWLEDGEMENTS

We thank the brain donors and their families for making our research possible. We also thank UCI neuropathologists and technicians at the Experimental Tissue Resource and ADRC for their assistance in procuring and imaging specimens, as well as for managing patient metadata. Tissue used to extend our analysis of associations with diabetic ketoacidosis was provided by the Human Brain Collection Core, Intramural Research Program, NIMH (http://www.nimh.nih.gov/hbcc) with the assistance of Dr. Stefano Marenco and Dr. Pavan Auluck.

---

**Fig. S1.**
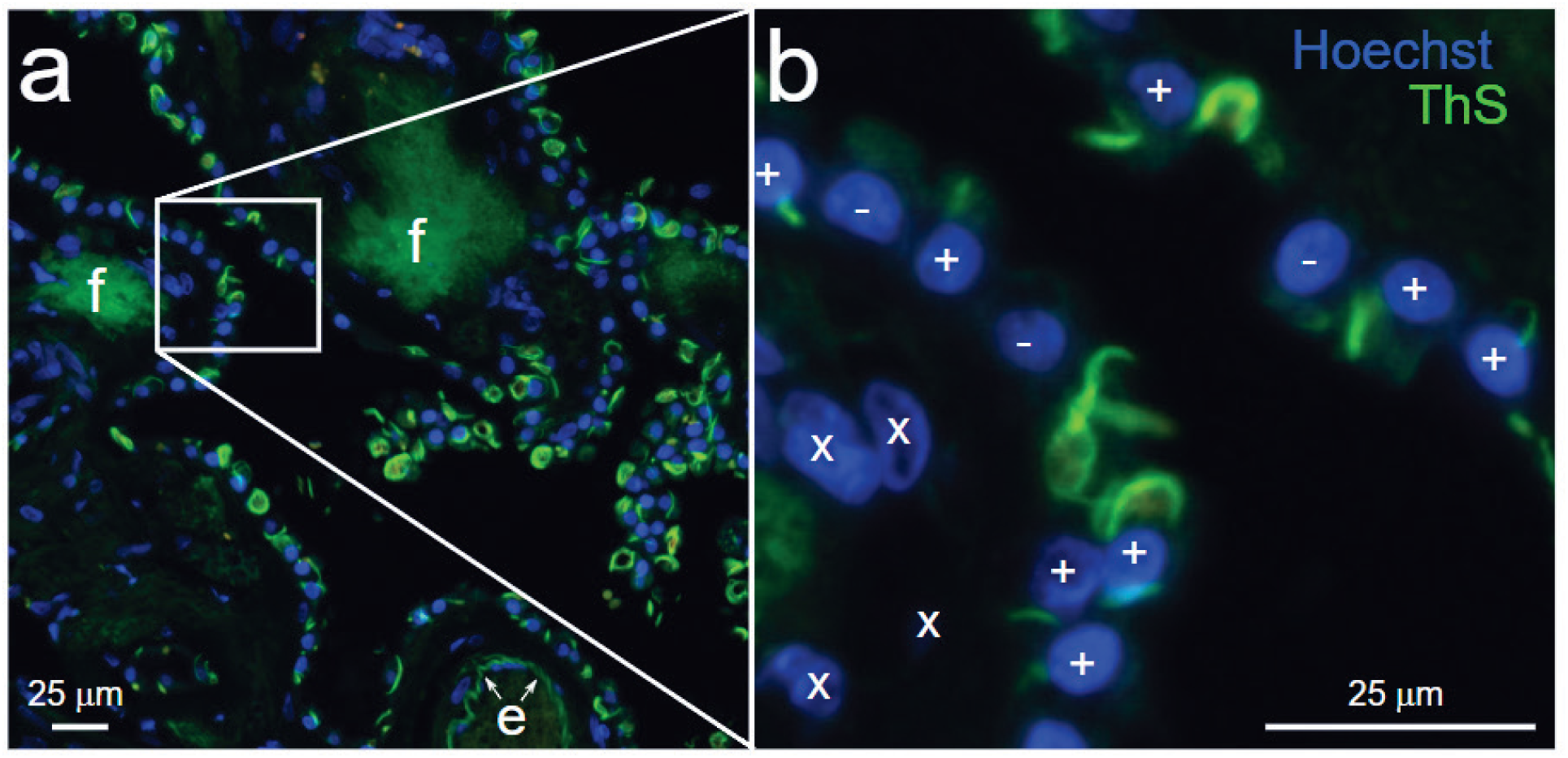
Manual annotations of BBs.**a**) An image tile used for manual annotations of ThS-positive and negative CPECs also may contain green-fluorescent features such as fibrosis (**f**) and elastin (**e**) that are ignored by annotators. **b**) Inset shows examples of CPECs that would be classified with respect to their containing any BB. Only CPECs with Hoechst-stained nuclei were annotated for BB presence (**+**) or absence (**-**). Hoechst-positive, non-CPEC cells in stroma (**x**) were not annotated.

**Fig. S2.**
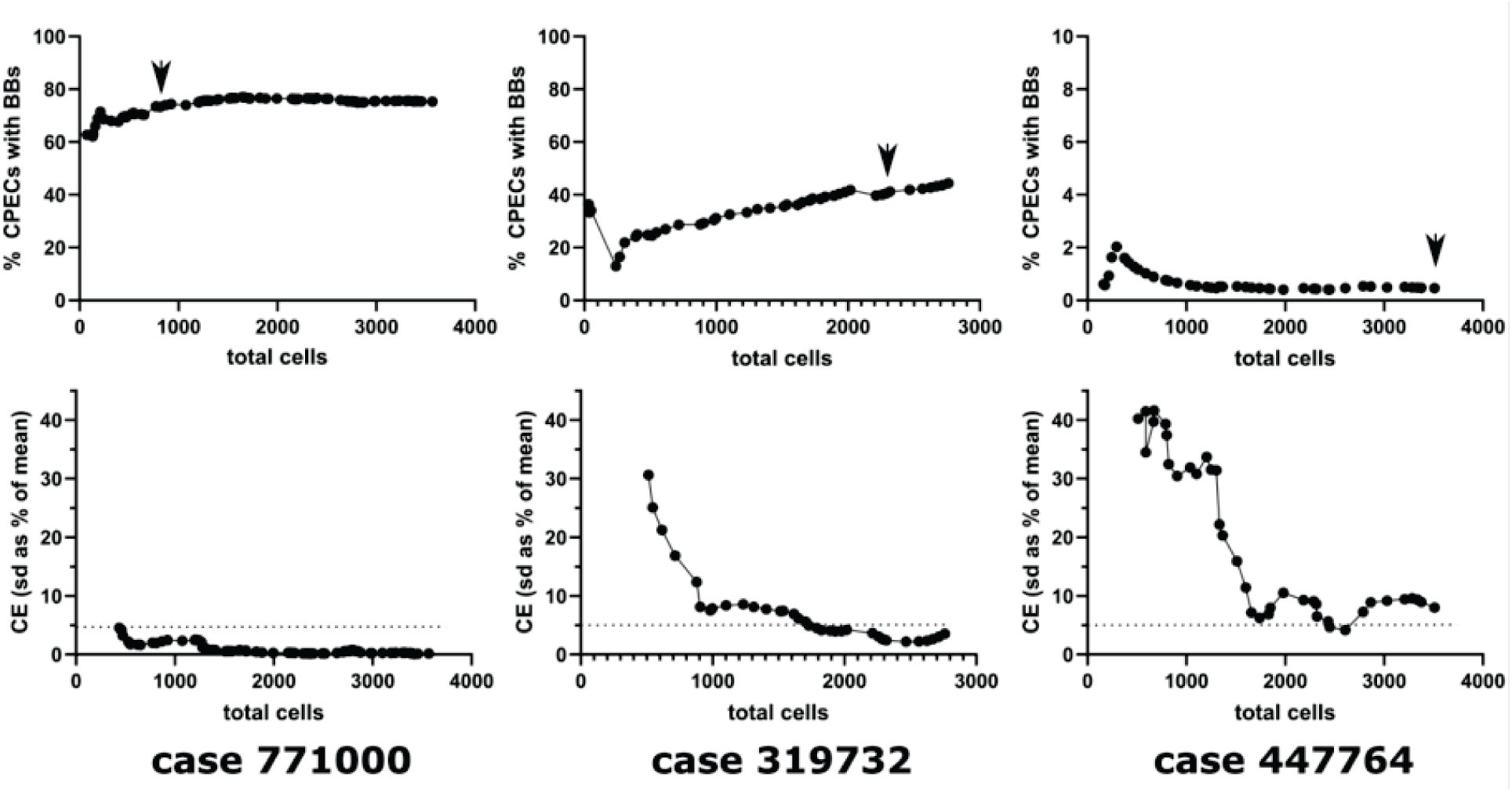
Criteria for completing manual annotation of a whole slide image. Image tiles cropped from a whole slide image (see **Fig.S1**) are presented randomly to annotators, who mark every Hoechst-positive CPEC in the tile. Upon completion of a tile, a cumulative estimate of the percentage of CPECs with ThS-positive BBs is updated (**upper panels**). Beginning at the tenth tile for which any CPECs were detected, the standard deviation of the estimate over the last ten tiles containing CPECs is calculated and expressed as a percentage of the mean of the last estimate, which generates a coefficient of error (CE) (**lower panels**). When ten consecutive tiles yield a CE of less than 5%, annotation of that whole slide image is considered complete (**arrowheads** for cases 771000 and 319732). If the estimate is that fewer than 2% of CPECs contain BBs after 3500 CPECs have been annotated, then annotation is considered complete after the tile containing the 3500th CPEC is annotated (case 447764).

**Fig. S3.**
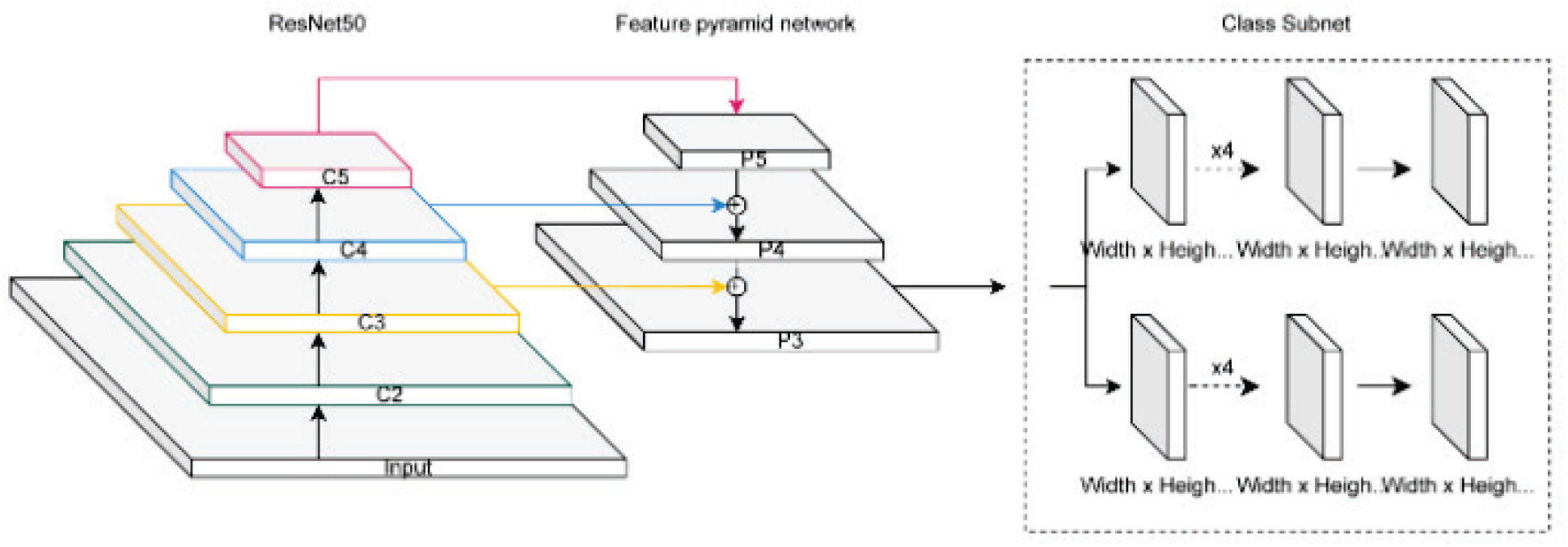
Modified RetinaNet model architecture used for CPEC detection. The architecture diagram shows the general structure used in our CPEC detection models. The main difference from the originally published architectures [27] is that only the P3 layer of the feature pyramid network is passed on to the class and box subnets.

**Fig. S4.**
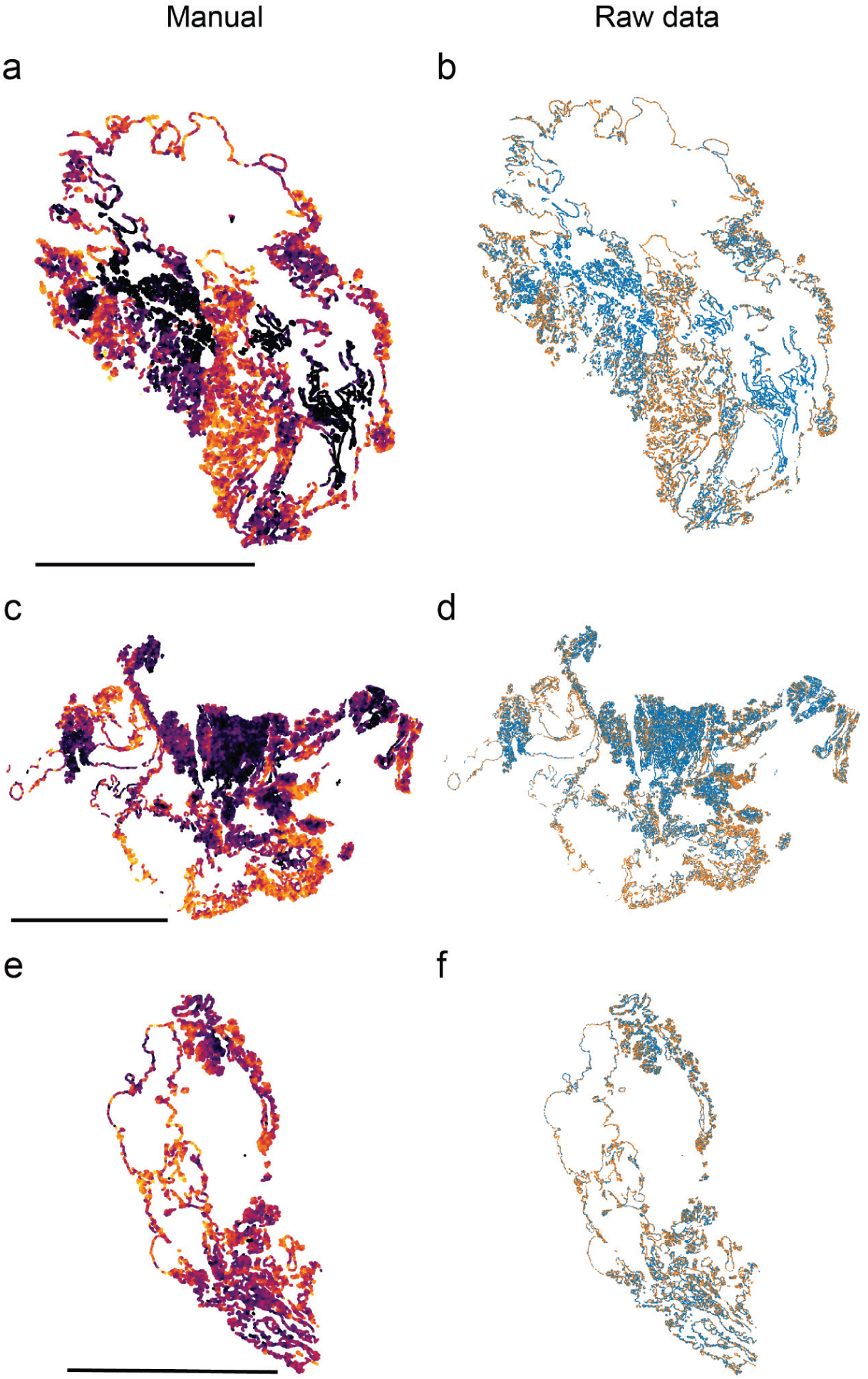
Transformation of spatial distributions into local densities. Every CPEC in whole slide images of ChP from three cases was annotated for the presence (orange symbol) or absence (cyan symbol) of BBs (right panels). These data then were transformed by calculating the percentage of CPECs within 100-μm of each CPEC that were BB-positive. These local densities were then represented as a heatmap-coded symbol at the position of each CPEC (right panels). Scale bars indicate 1 cm.

**Fig. S5.**
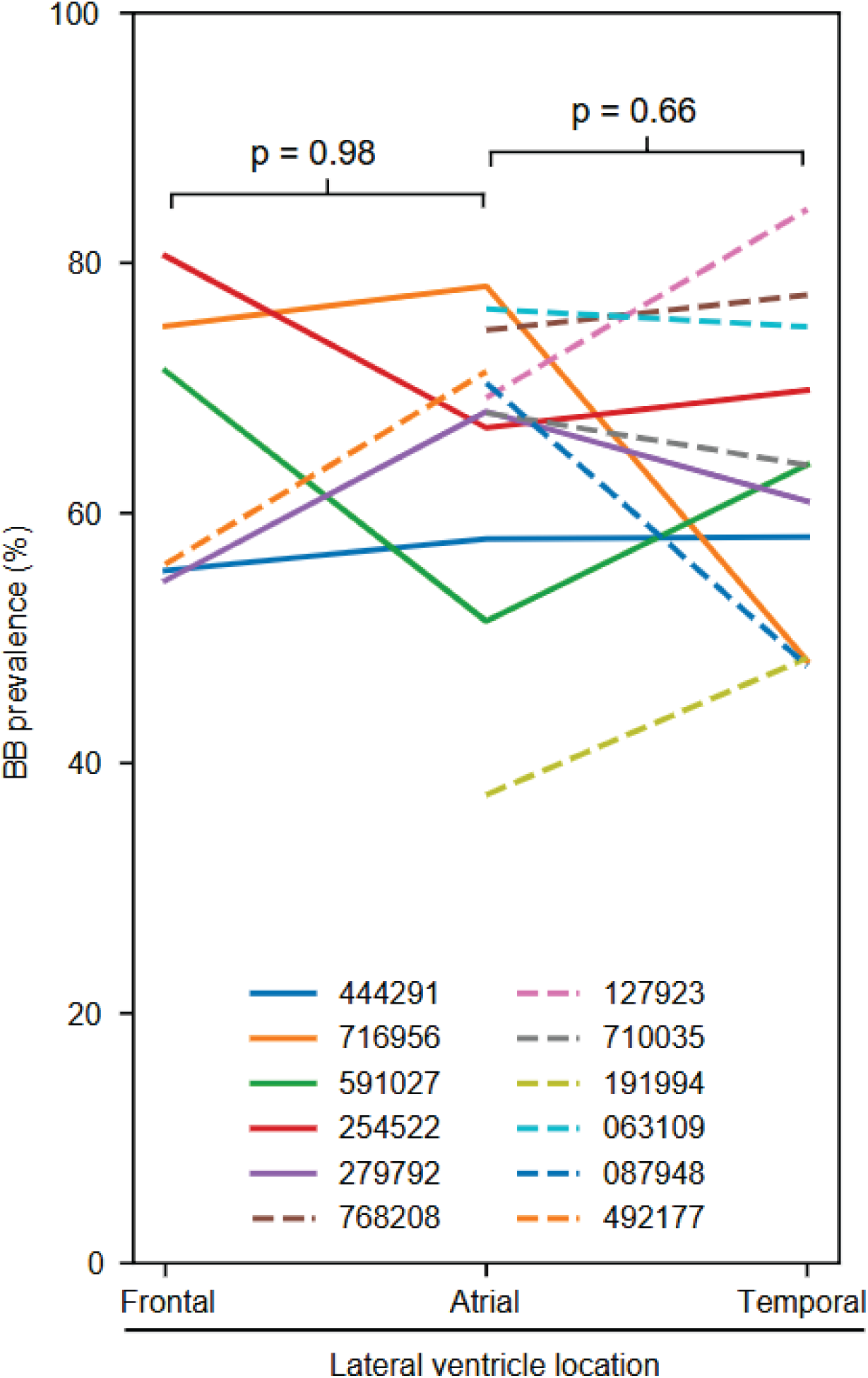
There is no systematic difference in the percentage of CPECs containing BBs among ChP samples taken from different locations within the lateral ventricle. P values are from paired t-tests for different locations from the same cases.

**Fig. S6.**
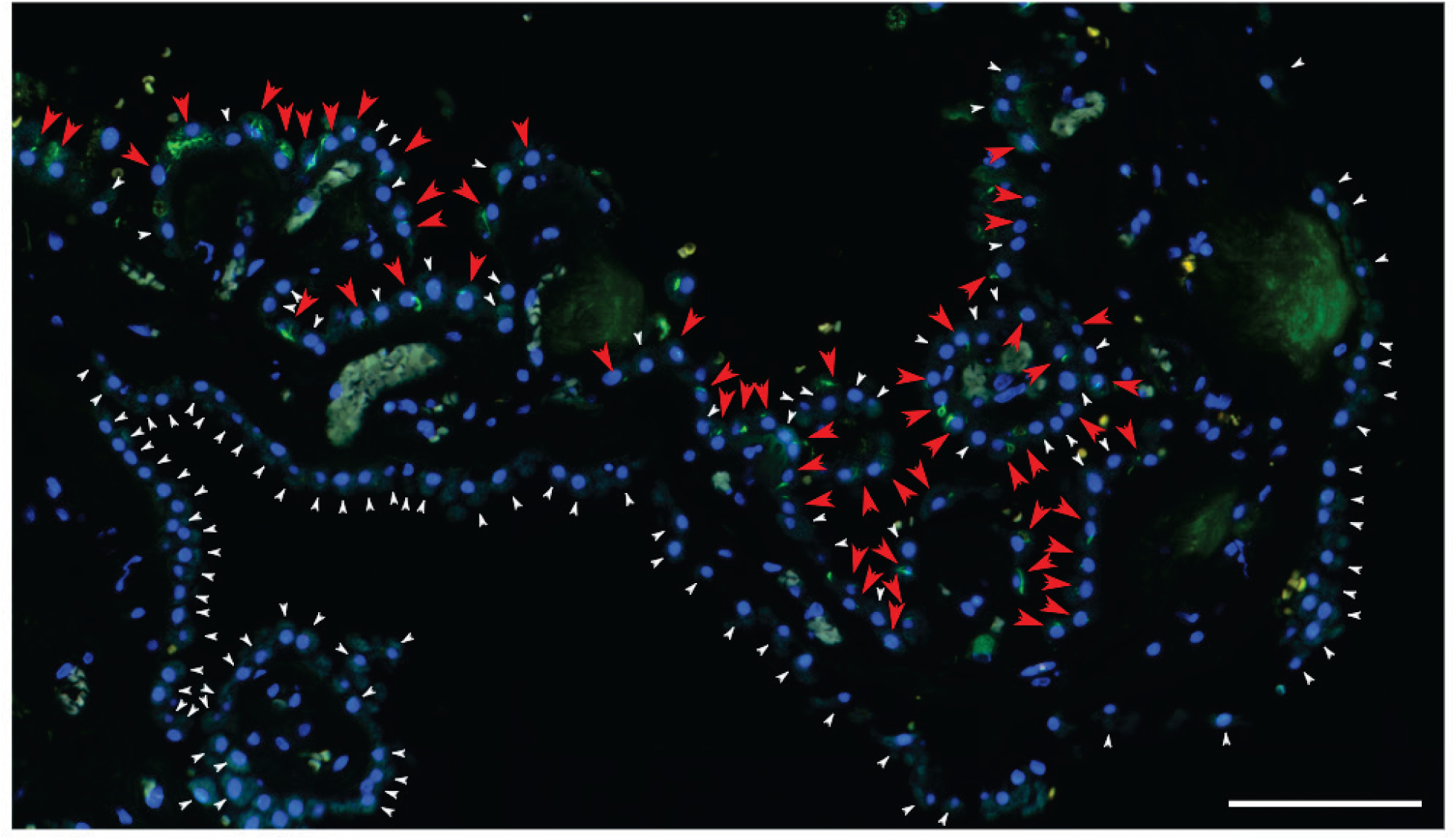
Clusters of BB-positive CPECs can be separated from regions of unaffected CPECs by thin stretches of stroma (case 363485). **Red arrowheads** indicate CPECs with BBs, while **white arrowheads** denote unaffected CPECs. Scale bar = 100 microns.

**Fig. S7.**
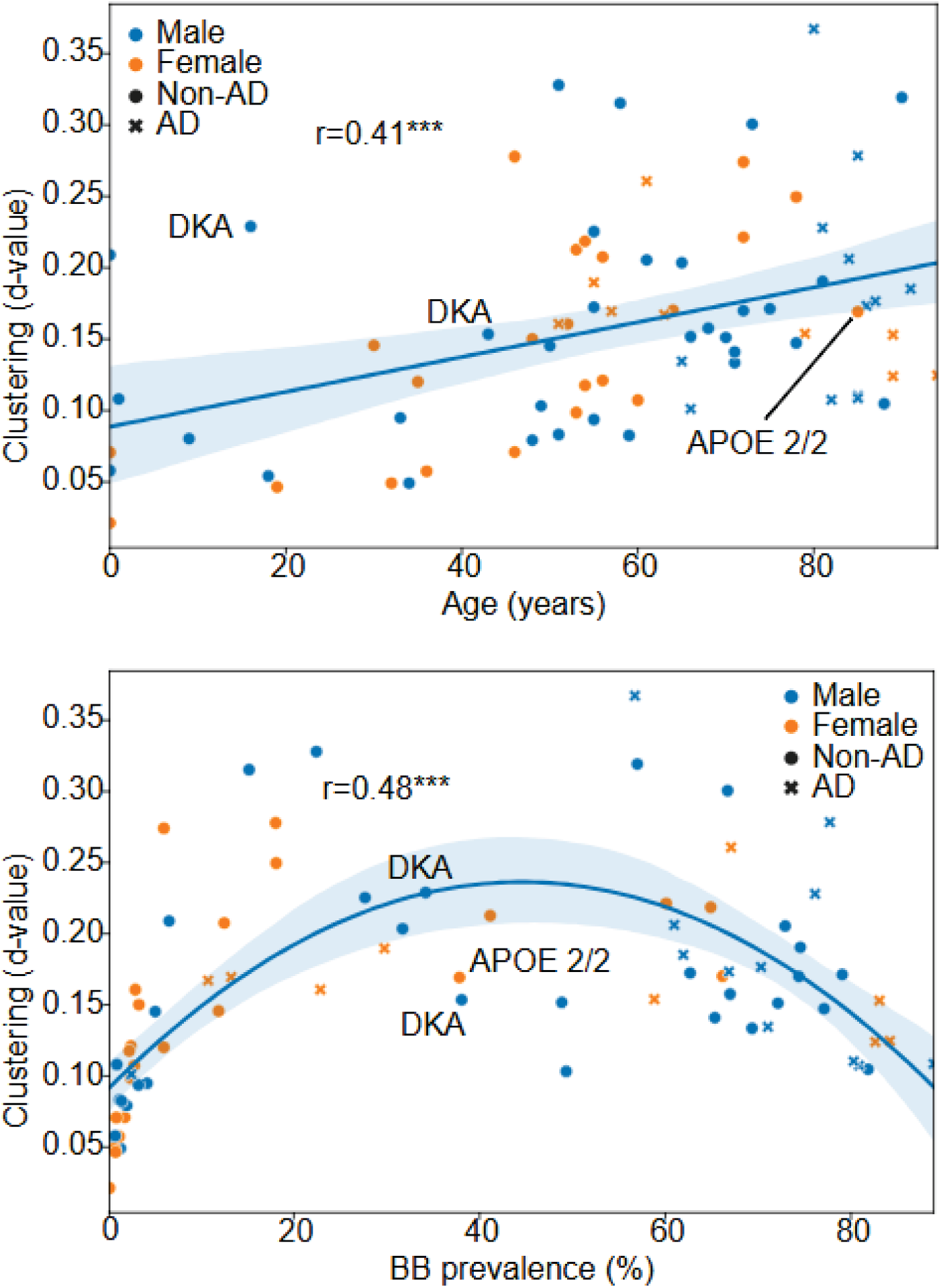
BB spatial clustering is associated with age and prevalence. Scatterplot showing DL-generated d-values from Kolmogorov–Smirnov statistical tests (a measure of the degree of clustering) as a function of age (**top**) and BB prevalence (**bottom**). Trendlines are linear (**top**) and 2^nd^ order polynomial (**bottom**) regressions with 95% confidence intervals (**blue shaded areas**). R-values are Spearman correlation coefficients. **Asterisks** indicate p < 0.001.

**Fig. S8.**
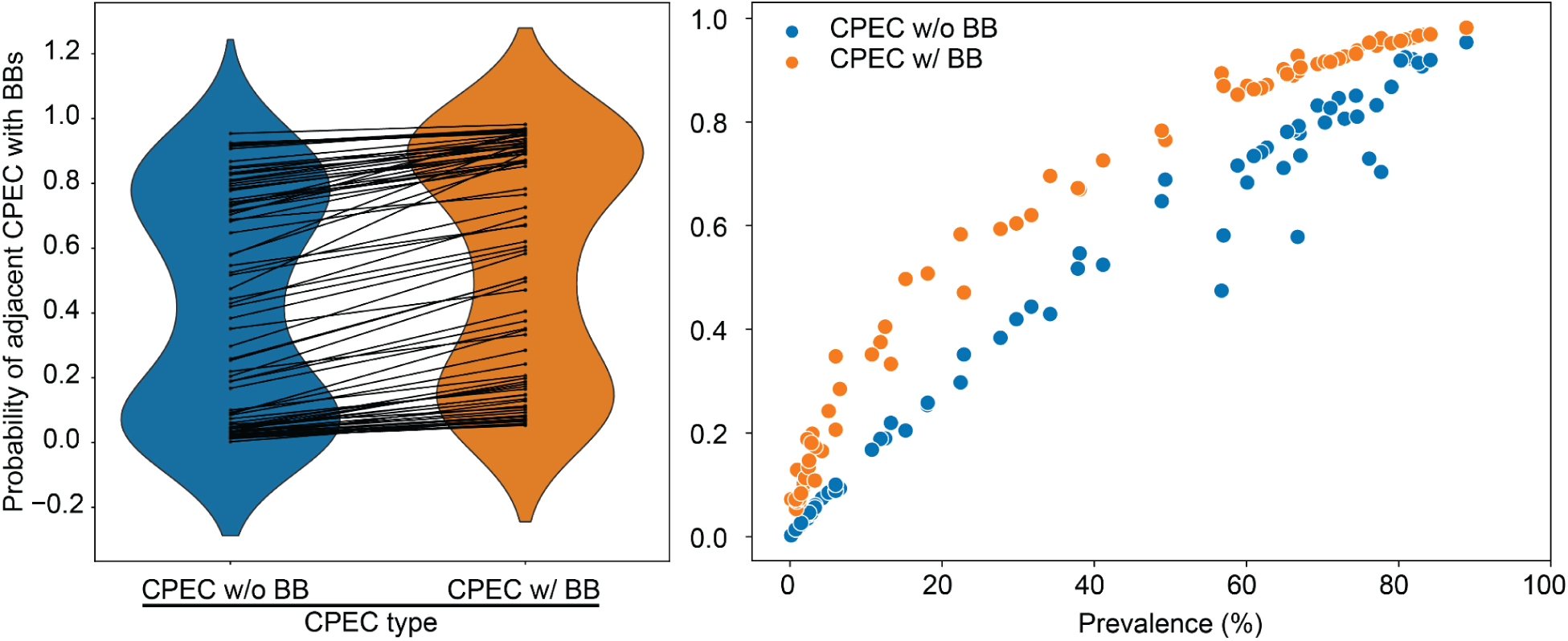
Similar to the analysis shown in Fig.8, BB+ CPECs are also more likely to be one CPEC removed from other BB+ CPECs. **Left**) Violin plots comparing the probability of a BB+ CPEC being one CPEC removed from a CPEC with or without a BB across the different cases. Lines between the violin plots connect data from the same case. **Right**) Scatterplot showing the probabilities previously mentioned as a function of BB prevalence.

